# Identification of Potential Dual-Targets Anti-Toxoplasma gondii Compounds Through Structure-Based Virtual Screening and In-Vitro Studies

**DOI:** 10.1101/827774

**Authors:** Nurul Hanim Salin, Rahmah Noordin, Belal Omar Al Najjar, Ezatul Ezleen Kamarulzaman, Muhammad Hafiznur Yunus, Izzati Zahidah Abdul Karim, Nurul Nadieya Mohd Nasim, Iffah Izzati Zakaria, Habibah A. Wahab

## Abstract

*Toxoplasma gondii* is the etiologic agent of toxoplasmosis, a disease which can lead to morbidity and mortality of the fetus and immunocompromised individuals. Due to the limited effectiveness or side effects of existing drugs, the search for better drug can didates is still ongoing. In this study, we performed structure-based screening of potential dual-targets inhibitors of active sites of *T. gondii* drug targets such as uracil phosphoribosyltransferase (UPRTase) and adenosine kinase (AK). First screening of virtual compounds from the National Cancer Institute (NCI) was performed *via* molecular docking. Subsequently, the hit compounds were tested *in-vitro* for anti-*T. gondii* effect using cell viability assay with Vero cells as host to determine cytotoxicity effects and drug selectivities. Clindamycin, as positive control, showed a selectivity index (SI) of 10.9, thus compounds with SI > 10.9 specifically target *T. gondii* proliferation with no significant effect on the host cells. Good anti-*T. gondii* effects were observed with NSC77468 (7-ethoxy-4-methyl-6,7-dihydro-5H-thiopyrano[2,3-d] pyrimidin-2-amine) which showed SI values of 25. This study showed that *in-silico* selection can serve as an effective way to discover potentially potent and selective compounds against *T. gondii*.

## Introduction

The parasite *Toxoplasma gondii* is an obligate intracellular protozoan that can infect all warm-blooded animals including human. It infects ∼30% of the human population worldwide, and the prevalence is up to 90% in some European populations [1, 2]. In Malaysia, chronic *Toxoplasma* infection was reported to vary from 10% to 50% [3]. In healthy immunocompetent adults, most infections are asymptomatic, but may cause lymphadenopathy and self-limiting flu-like symptoms. However, infection in immunocompromised patients such as those with AIDS can lead to fatal encephalitis. Furthermore, infection in seronegative pregnant women may lead to fetal infection that result in stillbirth, ocular diseases and mental retardations [1, 4, 5].

The most effective treatment for *T. gondii* infection is the synergistic combination of pyrimethamine and sulphonamides, especially sulphadiazine. Sulphonamides inhibit dihydrofolic acid synthetase while pyrimethamine inhibits dihydrofolate reductase, key enzymes in the synthesis of purines [1]. Although the treatment is quite effective, it has adverse effects especially in immunocompromised patients, such as haematological toxicity caused by pyrimethamine; and cutaneous rash, leukopenia and thrombocytopenia caused by the sulphonamide. The problems are compounded with the fact that the drug has no efficacy against tissue cysts, thus may encourage the appearance of resistant strains leading to the chance of recurrence after treatment [1, 4, 5]. Other drugs being used for toxoplasmosis include clindamycin, atovaquone and spiramycin [6]. Clindamycin is reported to be effective, but may cause ulcerative colitis [7]. Spiramycin has lesser anti-*T. gondii* effects compared to sulfadiazine and pyrimethamine; and it produces high tissue concentrations, particularly in the placenta, but without crossing the placental barrier. Thus, the development of new alternative drugs for the treatment of *T. gondii* infection is still needed to address the above limitations.

This study used two reported key targets of *T. gondii* namely uracil phosphoribosyltransferase (UPRTase) and adenosine kinase (AK). The pyrimidine and purine salvage pathways of *T. gondii* are good targets as these pathways are highly dissimilar between the parasite and host. In *T. gondii*, the parasite synthesizes pyrimidine nucleotide *de novo* and UPRTase is the only operative enzyme that salvages pre-formed pyrimidines to the nucleotide level [2]. Meanwhile, targeting AK is important because *T. gondii* needs to recover purine from the adenosine kinase since the parasite is unable to synthesize purine [8]. It is therefore interesting to attain a single compound which may be able to inhibit these two targets at once and potentially serve as a good lead for further downstream investigations.

In this study, virtual screening of anti-*T. gondii* was used to screen for potential drug candidates that have good complementarity in terms of shape and physico-chemical properties with the selected drug targets. The National Cancer Institute (NCI), USA database was used to virtually screen for anti-*T. gondii*. Subsequently pure compounds of the *in-silico* hits were then obtained from the NCI for *in vitro* validation. The selected pure compounds from the NCI database were tested using cell viability assay with Vero cells as host to determine cytotoxicity effects and drug selectivity.

## Methods

### Molecular docking

The protein-ligand docking using Autodock 3.05 for virtual screening workflow can be divided into three steps: the preparation of receptors and ligands structures, docking, and scoring [9]. The receptor targets selected were Uracil Phosphoribosyltransferase (UPRTase, PDB ID: 1UPF) and Adenosine Kinase (AK, 1LIJ) of *T. gondii*. In the receptor structure’s file preparation, the bound ligand and water molecules were removed from the crystal structure. Polar hydrogens were added followed by adding Kollman charges to the receptor’s amino acid and the partial charges were computed. The charged proteins were then solvated using the AutoDock3.0 “addsol” module and atom types were assigned for the receptor using the AutoDockTools [10].

In compound’s library files, the files were downloaded from NCI that have hydrogen atoms added to each compound, and subsequently converted from sdf format to mol2 format using OpenBabel [11]. The mol2 file, were separated into individual files and renamed and converted to the AutoDock pdbq format to specify the rotatable bonds for the docking conformational searches. The Gasteiger charges and maximum number of rotatable bonds were applied to all the ligands. The grid parameter file was set-up using AutoDockTools to constrain the search area. The parameters from the gpf file were taken by AutoGrid and the grid maps were generated for use with AutoDock. Pre-calculated grid maps for each atom type present were required in the ligand being docked. The centre of the grid box was specified at in x, y and z dimensions,which was the centre point for 5-fluorouracil binding site of UPRTase and AK.

Docking of ligand to protein was carried out using the new empirical free energy function and the Lamarckian Genetic Algorithm parameters (LGA) and 100 runs per simulation. A standard protocol such as cluster tolerance was less than 1.0 Å, an initial population of 50 randomly placed individuals, a maximum number of 15 × 10^5^ energy evaluations, a maximum number of generations of 2.7 × 10^4^, a mutation rate of 0.02, a crossover rate of 0.80 and an elitism value of 1 was employed [10, 12]. After the completion of docking calculation, AutoDock provided output of the docked coordinates and free energy of binding in the docking log file (dlg). The information regarding the ligand pose, structural coordinates, binding free energies, and cluster sizes were extracted out of the dlg file. The structures were converted into pdb formats.

### Virtual screening

Prior to the screening, the 1855 of NCI compounds were filtered according the druglikeness property Lipinski’s rule of 5 [13] using Accelrys Discovery Studio Client 2.5 [14] and were minimized by MM+ force field using Hyperchem 7.0 [15]. Then, the compounds from NCI databases were screened using the molecular docking method as described above and evaluated their binding to the receptor targets. Compounds scored favourably in term of the free energy of binding, the best ligand conformations and interaction with the binding site’s residues were chosen as hit compounds that may potentially act as anti-*T. gondii* agents.

### *In-vitro* cell viability assay

#### NCI compounds

From the screening of NCI database, pure compounds NSC77468, NSC660301, NSC81462, NSC38968, NSC81213, NSC 108972, NSC153391 and NSC343344 were selected and obtained free of charge from the National Cancer Institute (NCI), Washington, DC, USA.

#### Positive controls

Clindamycin [CAS: 21462-39-5] and 5-fluorouracil [CAS: 51-21-8] were purchased from Sigma-Aldrich, St. Louis, MI, USA.

#### Host cells

The mammalian Vero cell line was obtained from the Institute for Research in Molecular Medicine (INFORMM), USM. The cell line originated from kidney of a normal adult African green monkey and was established in 1962 by Yasummura and Kawakita at the Chiba University, Japan (American Public Health Association, 1992).

*T. gondii* **RH strain maintenanc**e

*T. gondii* tachyzoites RH strain were obtained from the Institute for Research in Molecular Medicine (INFORMM), USM. The tachyzoites were maintained by passaging using intraperitoneal inoculation into healthy Swiss albino mice every 2-3 days. All experimental procedures that involved animals were performed according to USM ethical guidelines (USM/PPSF50 (003) JLD2) under internationally accepted principles for laboratory animal use and care.

#### Counting of cells

Vero cells were harvested by trypsinization when it reached 80-90% confluence and centrifuged at 1000 × *g* for 5 mins (Eppendorf Centrifuge 5804R, Germany). The clean Vero cell pellet was resuspended thoroughly with growth medium (GM) which comprised Dulbecco’s Modified Eagle medium [DMEM] (Gibco, USA) containing 0.5% penicillin-streptomycin (PS) and 5% fetal bovine serum (FBS). The number of cells was counted with a hemocytometer under inverted light microscope, using the trypan blue dye-exclusion method.

#### Counting of parasites

*T. gondii* was freshly harvested from mice peritoneal fluid and collected in a tube. The tube was centrifuged at 200 × *g* for 10 mins. The resulting supernatant which contained more than 90% pure tachyzoites was collected; while the pellet which contained mouse macrophage cells was discarded. The supernatant was centrifuged again at 1000 × g for 10 mins to pellet the tachyzoites. The clean *T. gondii* pellet was resuspended thoroughly with wash medium (WM) which comprised DMEM with 0.5% PS and without FBS; and the number of viable *T. gondii* was counted as above.

#### MTS and PMS reagent

The detection reagents composed of solutions of the tetrazolium compound MTS (Promega Madison, WI, USA) catalog number G1112 and an electron coupling reagent (PMS) [CAS: 299-11-6] from Sigma (St. Louis, MO,USA).

#### Cytotoxicity effect on Vero cells

Vero cells were harvested by trypsinization when it reached 80-90% confluence. A volume of 100 *µ*l Vero cells (6 *×* 10^4^ cells/ml) in GM was cultured in triplicate wells of a 96-well plate and incubated for 24 hours. Then 100 *µ*l of GM was added into each well, which contained different concentrations of the NCI compounds and clindamycin, from 1.5625 to 100 *µ*g/ml. The negative control well contained 100 *µ*l Vero cells and phosphate buffered saline, pH7.2 (PBS) in control wells for NCI compounds and clindamycin. At each concentration, the NCI compounds and clindamycin were assayed in triplicates and incubated for 24 hours [16–18].

After treatment for 24 hours, the anti-*T. gondii* activity was examined using MTS-PMS assay. MTS solution was prepared (2 mg/ml) by dissolving the powder in PBS. Exposure to light was avoided during the preparation. PMS (0.92 mg/ml) was similarly prepared in PBS and both solutions were stored at −20° C. A volume of 20 *µ*l of MTS solution was added into each well and the cells were incubated for one hour in a CO_2_ incubator [19, 20]. The absorbance (abs) was then measured at 490 nm using a microplate reader (Thermo Scientific Multiskan^®^ Spectrum, USA). The percentage of cell viability was calculated using the following formula [21, 22].

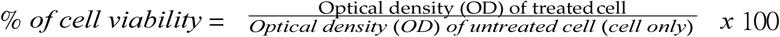

#### Treatment of *T. gondii*-infected Vero cells

Vero cells were harvested by trypsinization when it reached 80-90% confluence. A volume of 100 *µ*l diluted Vero cells of (6 × 10^4^ cells/ml) in GM was cultured in triplicate wells of a 96-well plate and incubated for 24 hours. Then, 100 *µ*l of tachyzoites (3 *×* 10^5^ /ml) in GM2 (comprising DMEM, 0.5 % PS and 2% FBS) were added into the Vero cell-seeded wells. A volume of 100 ul the above medium was added to the negative control well and the plate was incubated for six hours. Subsequently, the wells were washed twice with WM to remove non-adherent parasites. A volume of 100 *µ*l of GM2 was added and again incubated for 18 hours. A volume of 100 *µ*l of GM2 was added to each well with different concentrations of the NCI compounds and clindamycin as positive control (from 0 to 25 g/ml; rows A to G respectively). The negative control well contained100 *µ*l Vero cells and PBS in control wells for NCI compounds. At each concentra tion, the NCI compounds and clindamycin were assayed in triplicates and incubated for 24 hours. After treatment for 24 hours, anti-*T. gondii* activity was examined using MTS-PMS assay and the results analysed using the same formula as above.

The percent cell viability was proportional to the mortality of *T. gondii*. The graphs were plotted using Prism 5.0 (GraphPad Software Inc., CA, USA). The median effective concentration (EC_50_) value refers to the concentration of the compounds/clindamycin necessary to inhibit 50% of the control values. Selectivity refers to the mean of the EC_50_ value for Vero cells relative to the mean of the EC_50_ value of the *T. gondii*. A P value of less than 0.05 was considered statistically significant and all data points represent the mean of three independent experiments. SI (selectivity index) measures the strength of inhibition against the Vero cells and the *T. gondii*. High SI indicates that the compound specifically targets the proliferation of *T. gondii* with negligible effect on Vero cells [16, 23, 24].

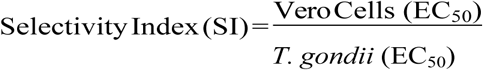

## Results and discussion

### Control Docking

Prior to the virtual screening, control docking was carried out on two *T. gondii* targets namely UPRTase and AK against its known inhibitors. The crystal conformation of UPRTase (PDB ID: 1UPF) and 5-fluorouracil were reproduced with good root-mean-square deviation (RMSD) value of 0.59 Å. The free energy of binding calculated was −4.22 kcal/mol and inhibition constant, K_*i*_ of 0.809 mM (Fig 1). Whilst, the docking of 2,7-iodotubercidin into AK (PDB ID: 1LIJ) also showed low RMSD (0.51 Å) with the free energy of binding of −10.99 kcal/mol and predicted inhibition constant, K_*i*_ of 8.87 nM (Fig 2).

**Fig 1.**
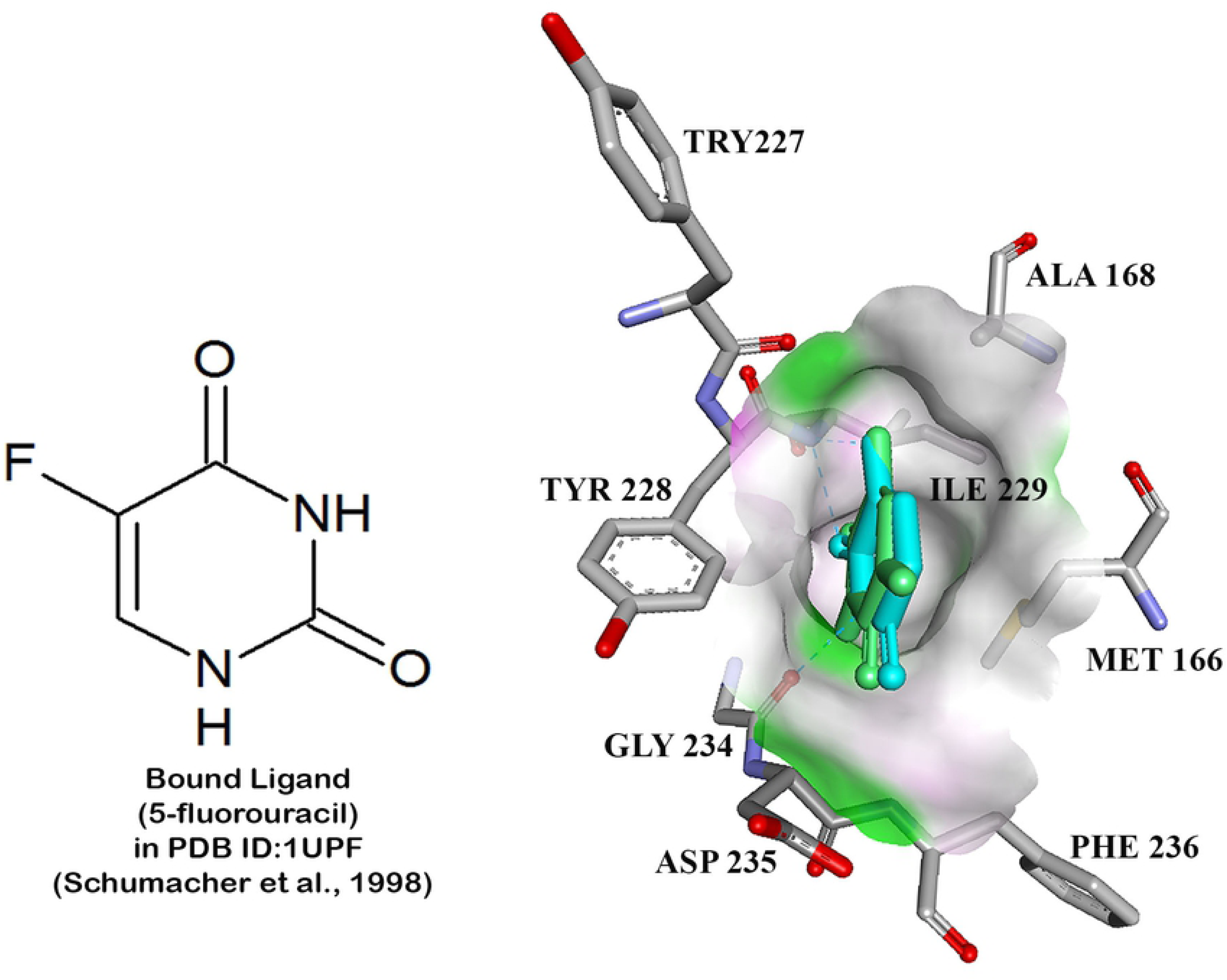
The superimposition of docked conformation of5-fluorouracil (green) and the conformation of crystal structure (blue) in the binding pocket of UPRTase for control docking

**Fig 2:**
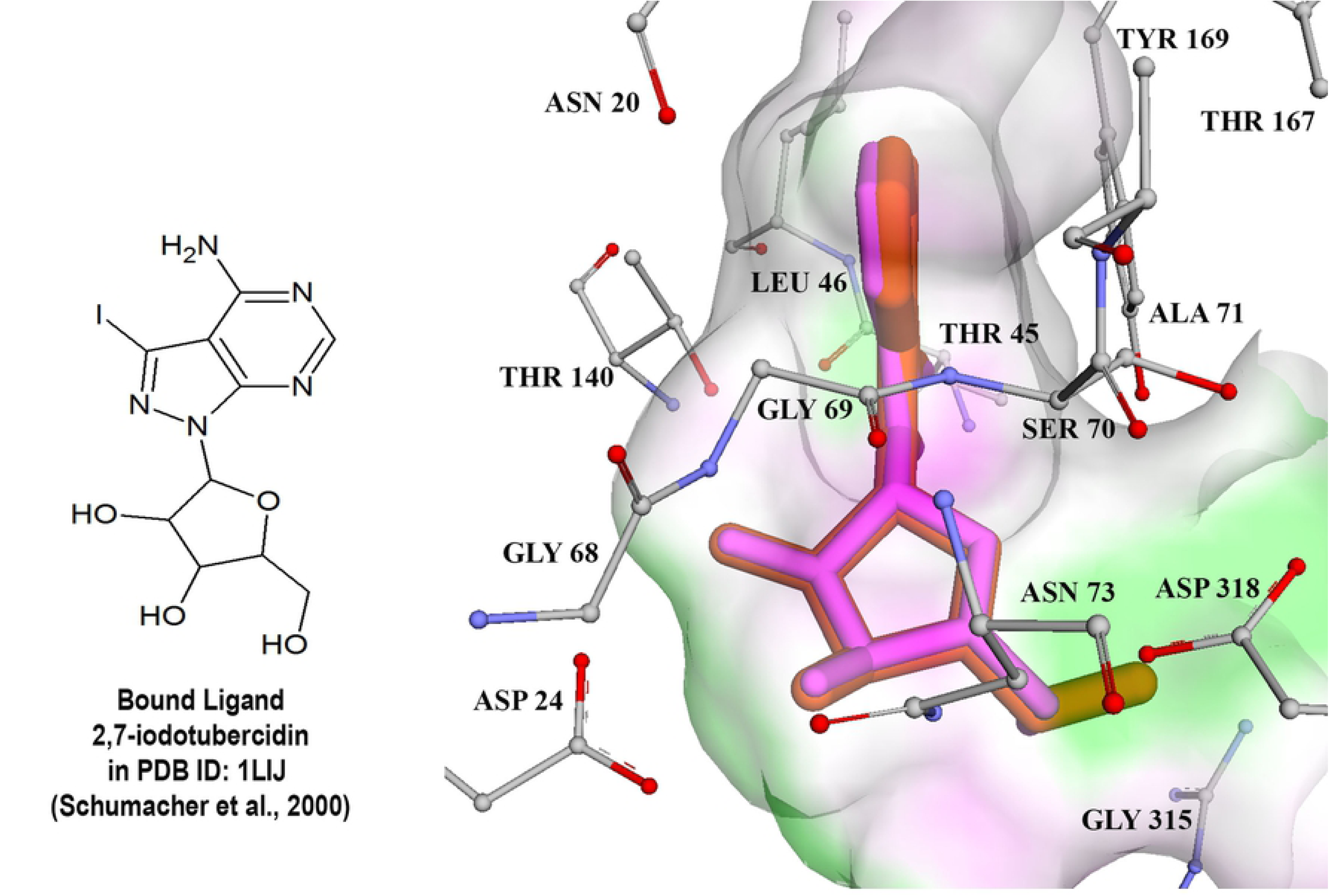
The superimposition of docked conformation of 2, 7-iodotubercidin (pink) and the conformation of crystal structure (orange) in the binding pocket of AK for control docking

The RSMD values obtained from this study were all below 1 Å with favourable free energy of binding, thus these docking parameters could be used for subsequent dockings (virtual screening). The best conformations and orientation of known inhibitors were generated using Lamarckian genetic algorithm in AutoDock 3.0.5 while keeping the enzyme structure rigid.

### Virtual screening of NCI dat a b a s e

Virtual screening was performed on 1216 drug-like compounds from the NCI database target ed to UPRTase and AK, using AutoDock 3.0.5. First, the top 25 compounds which bound favourably to at least two or more enzymes were selected based on the lowest free energy of binding and were considered for testing in biological assay. Then, the compounds were ranked according to the best ligand conformations. Eight of the 25 top ranked NCI compounds, i.e. NSC77468 (7-ethoxy-4-methyl-6,7-dihydro-5H-thiopyrano[2,3-d]pyrimidin-2-amine), NSC660301(1-(2-amino-4-chlorophenyl)pyrrolidine-2,5-dione), NSC81462 (dodecahydro-1,4,9b-triazaphenalene), NSC38968(2-(piperazin-1-yl)ethan-1-amine), NSC81213(4-chloro-2-nitrobenzene-1-sulfonamide), NSC108972(3-methyl-1,1-dioxo-1 H-1-benzothiophene-2-carboxamide), NSC153391(3-[(3-aminophenyl)sulfanyl]-1-(2-methoxyphenyl)pyrrolidine-2,5-dione) and NSC343344 (6-aminopyridin-3-yl)(thiophen-2-yl)methanone) (Fig 3)(Table 1) were selected for subsequent investigations. These top best compounds were analyzed for their binding interaction such as hydrogen bonding, hydrophobic interaction and aromatic interaction in order to further characterize their mechanism of action in the active site.

**Table 1.** Eight top NCI compounds with favourable binding properties to UPRTase and AK of *T. gondii*

| No | Compound Name | No. of ligand Conformation | FEB kcal/mol | Predicted Inhibition Constant, Ki, (μM) |
| --- | --- | --- | --- | --- |
| UPRTase |  |  |  |  |
| 1 | NSC81462 | 79 | -9.26 | 0.163 |
| 2 | NSC81213 | 78 | -9.58 | 0.096 |
| 3 | NSC660301 | 67 | -8.23 | 0.922 |
| 4 | NSC77468 | 19 | -7.73 | 2.140 |
| 5 | NSC38968 | 18 | -8.04 | 1.290 |
| 6 | NSC153391 | 18 | -7.67 | 2.390 |
| 7 | NSC108972 | 17 | -9.66 | 0.083 |
| 8 | NSC343344 | 4 | -8.49 | 0.599 |
| AK |  |  |  |  |
| 1 | NSC77468 | 58 | -15.46 | 0.005 |
| 2 | NSC81462 | 48 | -16.02 | 0.002 |
| 3 | NSC343344 | 44 | -13.89 | 0.066 |
| 4 | NSC108972 | 30 | -13.81 | 0.075 |
| 5 | NSC38968 | 19 | -13.13 | 0.238 |
| 6 | NSC660301 | 13 | -14.16 | 0.042 |
| 7 | NSC81213 | 9 | -13.49 | 0.130 |
| 8 | NSC153391 | 5 | -12.43 | 0.774 |

**Fig 3.**
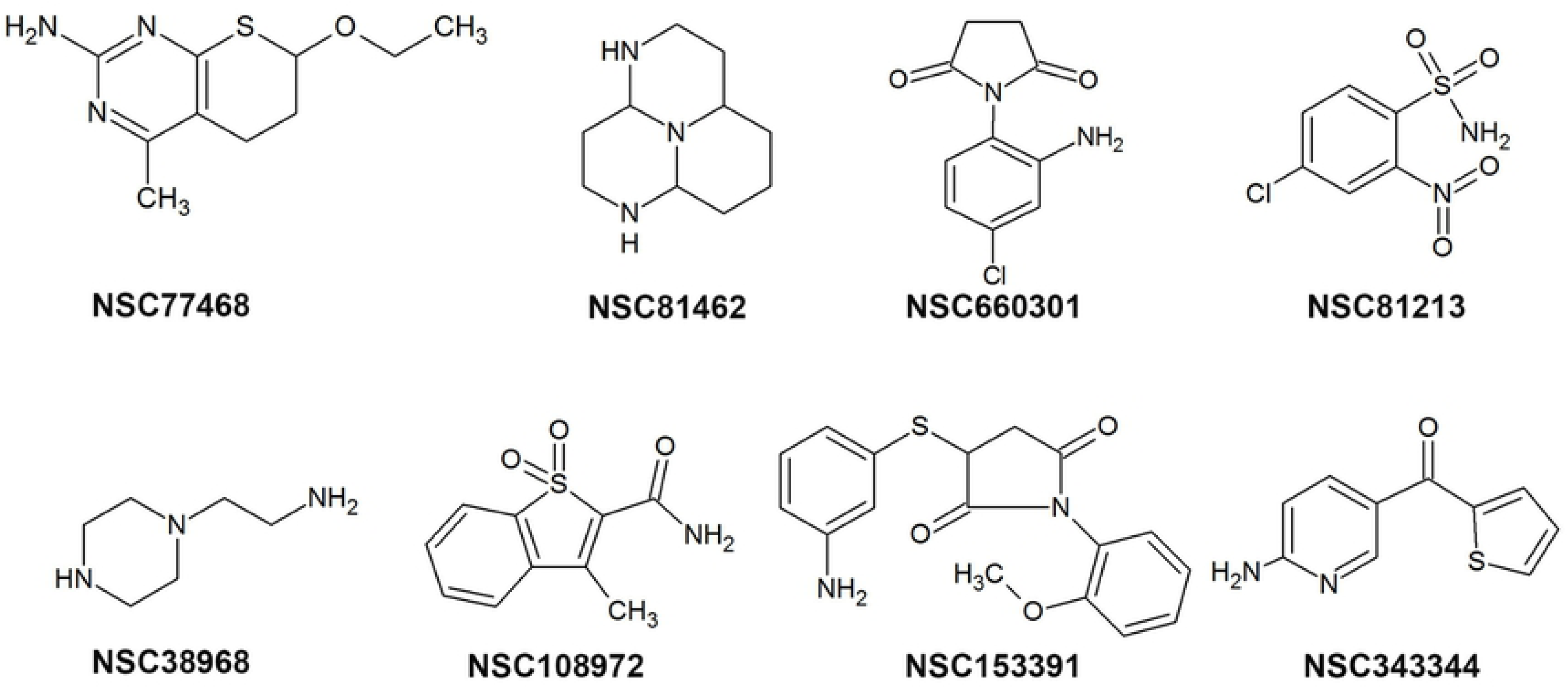
Top eight NCI compounds target to UPRTase and AK.

NSC77468 was found to have the best favourable hydrogen bond interaction (Fig 4) and significant number of hydrophobic interactions (Fig 5) with both the target enzymes i.e. UPRTase and AK. NSC77468 also was found to have the best aromatic interaction bound to UPRTase and AK (Fig 6). By inhibiting the parasite with multi-target inhibitors, the complementarities binding in getting the best interactions were also enhanced. Based on the analysis of the binding site interaction among the NCI compounds, NSC77468 was found as the most ideal candidate for anti-*T. gondii* compound among the eight top NCI compounds based on lowest free energy of binding, the best ligand conformations, and good intermolecular interactions with the targets.

**Fig 4.**
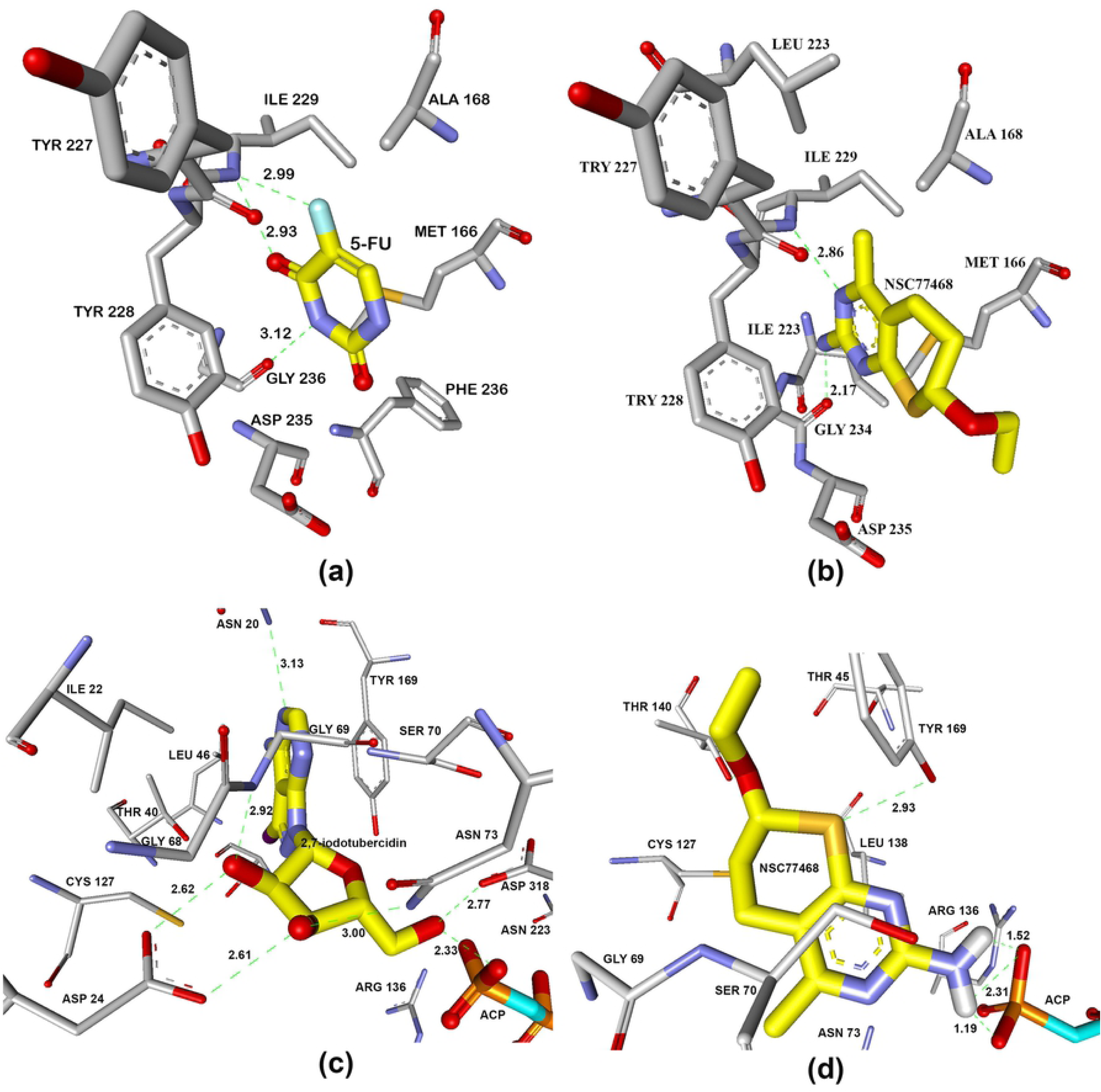
Hydrogen bond interaction formed between (a) 5-fluorouraciland (b) NSC77468 with ILE229, GLY234 in the active site of UPRTase. Hydrogen bond interaction formed between (c) 2,7-iodotubercidin (blue carbon) and (d) NSC77468(blue carbon) with amino acid residues (white carbon), ACP799 (yellow carbon)in AK active site. Diagram generated from Discovery Studio 4.1.

**Fig 5.**
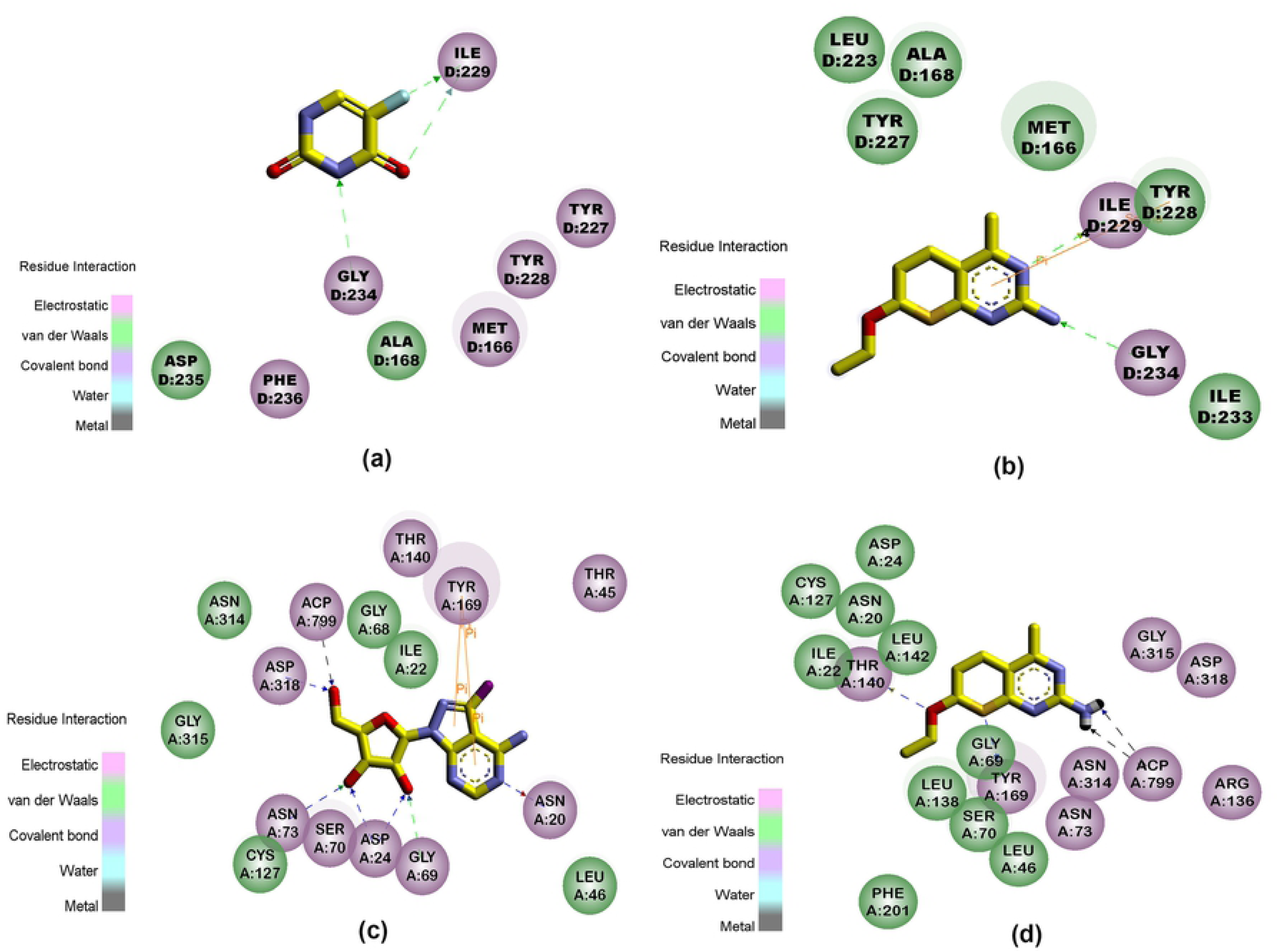
Interaction formed between 5-Fluorouracil, 2,7-iodotubercidin and NSC77468 with amino acid residues of (a)UPRTase and b) AK. Diagram generated from Discovery Studio 4.1; green residues represent non-polar contact while purple residues represent polar contact.

**Fig 6.**
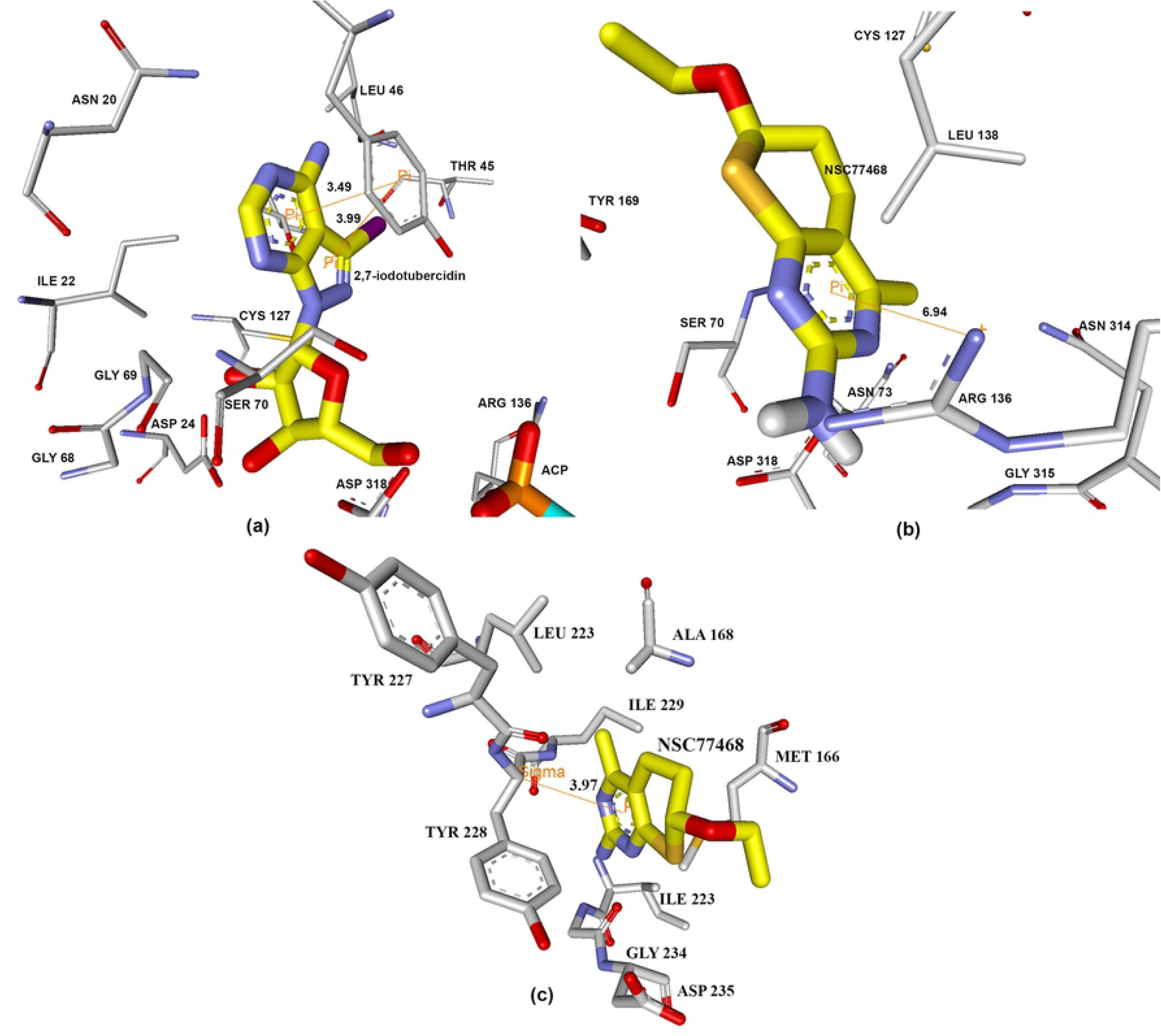
Aromatic interaction formed between (a) 2,7-iodotubercidin with TYR 169 in the active site of AK while NSC77468 with (b) ARG 136 and (c) TYR 228 in the active site of AK and UPRTase respectively.

### Structure-activity relationship

Based on the structure-activity relationship for the binding of ligands to UPRTase, of the eight NCI compounds only NSC77468 and NSC343344 can be considered as uracil analogue or pyrimidine analogue. The latter compound can be considered to be uracil analogues in which endocyclic imine or methylene group are replaced with endocyclic methylene or imine groups [25]. Table 2 show s the comparison of the substituent effects at the various positions pyrimidine ring of prodrug 5-fluorouracil and NSC77468 in the binding location of UPRTase. Based on the comparison, the differences were at C5-position for 5-fluorouracil and C6-position for NSC77468. Exocylic substituent of fluorine at 5-position may abolish the binding of 5-fluorouracil due to fluorine which has electron-withdrawing property. The exocyclic subtituents of flourine at 5-position will increase the level of ionization of this compound relative to the uracil [25]. The ionization occurs predominantly through the loss of N3 proton rather than the N1 proton [25]. Based on the binding of the NCI compounds at the binding site of AK, only NSC77468 showed hydrogen bond interaction with the substrate; a nonhydrolysable adenosine triphosphate analog (ACP) near Arg136. Guanidinium group at the side chain of Arg136 has a function to activate and stabilize the phos phate during catalysis (Fig 7). Therefore, based on this structure-activity relationship, NSC77468 could be developed into a lead compound which might provide better activity than 5-fluorouracil.

**Table 2.**
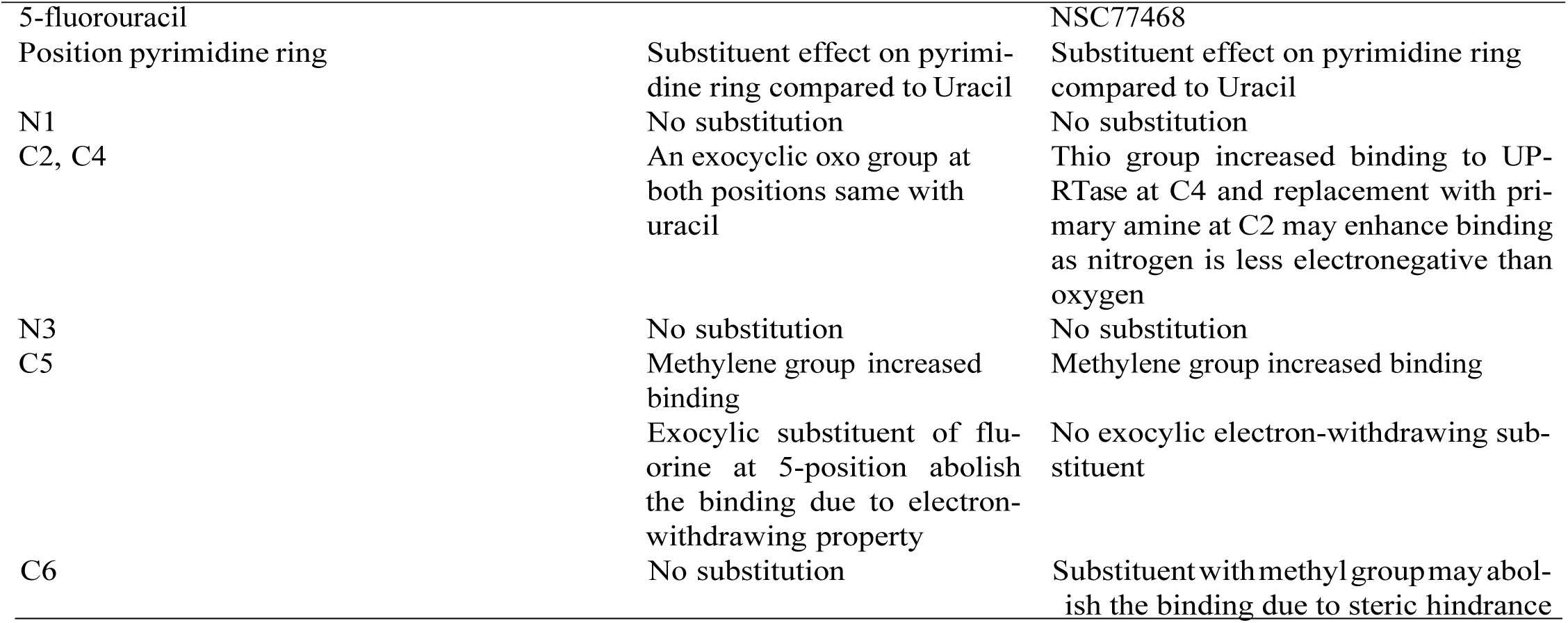
Structure-activity relationship for the binding of prodrug 5-fluorouracil and NSC77468 to UPRTase

| 5-fluorouracil<br>Position pyrimidine ring | Substituent effect on pyrimidine ring compared to Uracil | NSC77468<br>Substituent effect on pyrimidine ring compared to Uracil |
| --- | --- | --- |
| N1 | No substitution | No substitution |
| C2, C4 | An exocyclic oxo group at both positions same with uracil | Thio group increased binding to UPRTase at C4 and replacement with primary amine at C2 may enhance binding as nitrogen is less electronegative than oxygen |
| N3 | No substitution | No substitution |
| C5 | Methylene group increased binding | Methylene group increased binding |
|  | Exocyclic substituent of fluorine at 5-position abolish the binding due to electron-withdrawing property | No exocyclic electron-withdrawing substituent |
| C6 | No substitution | Substituent with methyl group may abolish the binding due to steric hindrance |

**Fig 7.**
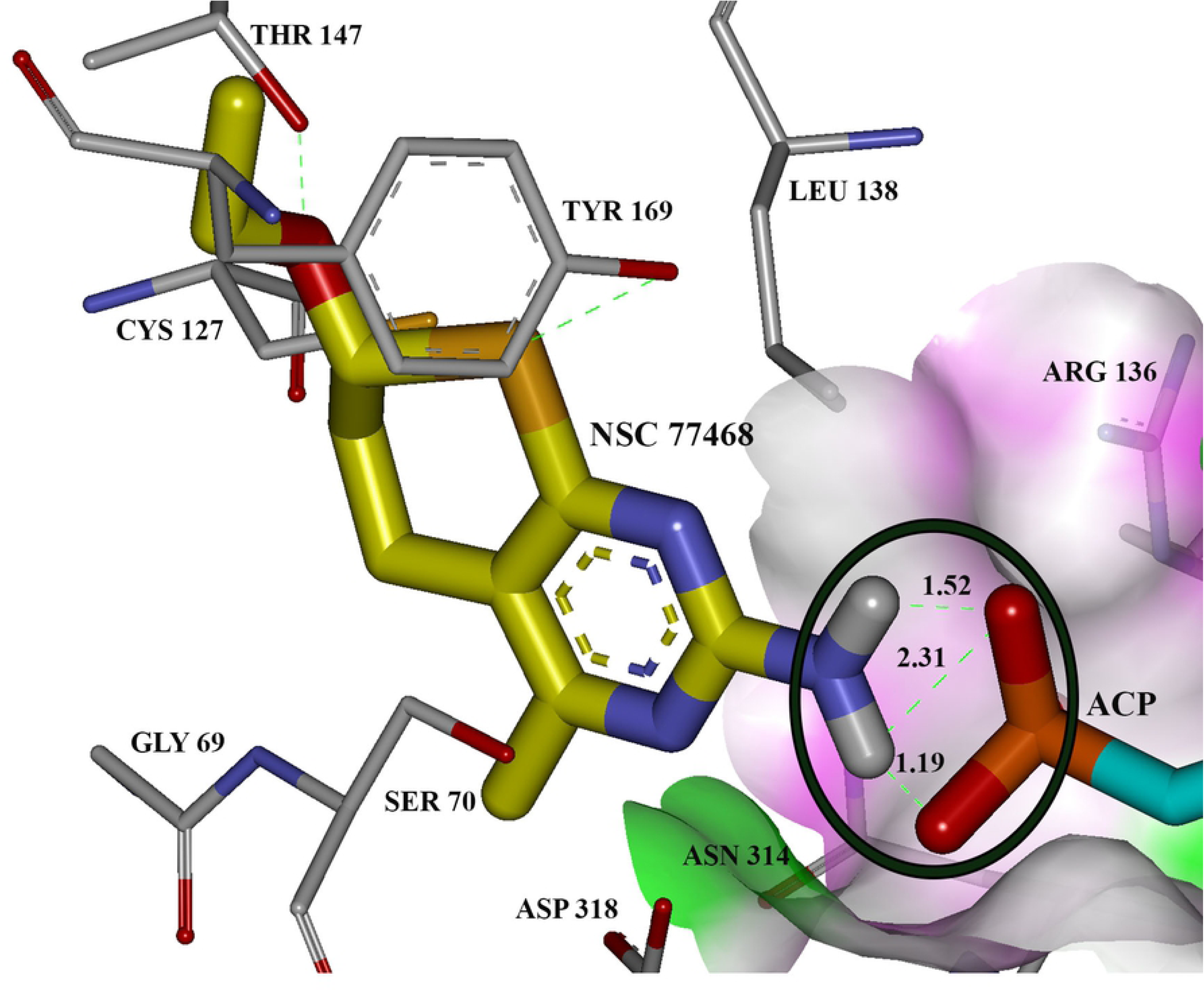
The hydrogen bond interaction of NSC77468 (yellow carbon) together with the substrate, ACP (blue carbon) in the binding location of AK.

### *In-vitro* cell viability assay

#### Positive Control

##### Clindamycin

From the *in-vitro* cytotoxicity assay, clindamycin showed EC_50_ of 46.63 *µ*g /ml against Vero cells (Fig 8). Meanwhile, the positive control drug, 5-flourouracil exhibited EC_50_ of 9.720 *µ*g/ml indicating high t oxicity against Vero cells. Compared to 5-flourouracil, clindamycin showed moderate toxicity and thus was used as the positive control in this study. From the anti-*T. gondii* MTS/PMS assay, clindamycin showed EC_50_ of 4.27 *µ*g/ml or 9.3 nM. The selectivity index (SI) of clindamycin was thus 10.9 (46.63 *µ*g/ml /4.270 *µ*g/ml) indicating that it has anti-*T. gondii* effect and low toxicity on the mammalian Vero cells.

**Fig 8.**
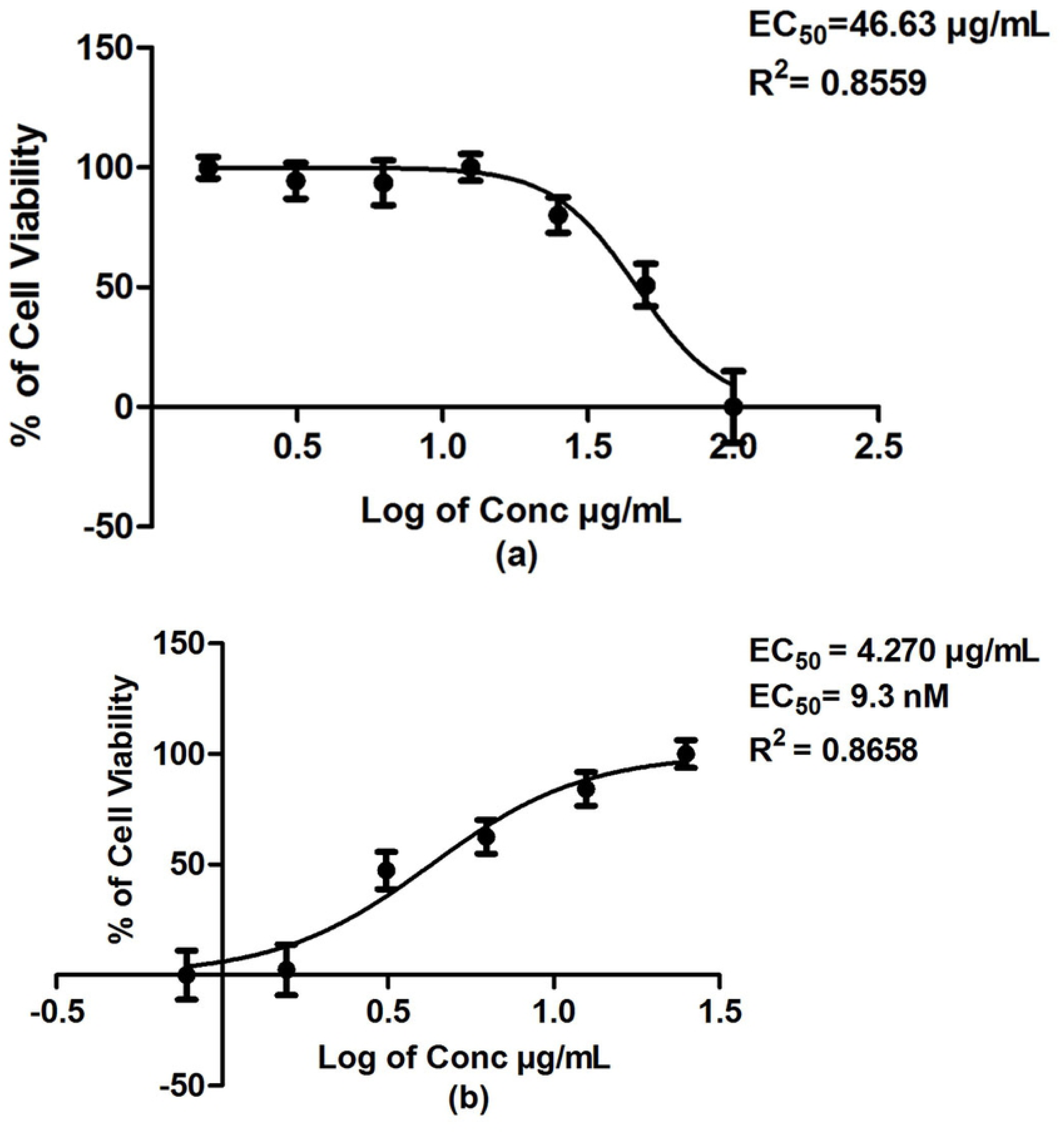
(a) Cytotoxic effect against Vero cells upon increasing clindamycin concentration; and (b) *T. gondii*-infected Vero cells treated with increasing clindamycin concentration.

##### Screening of NCI compounds

The aim of the bioassay in this study was to evaluate the efficacy and activity of the top few hit compounds in *in-vitro* culture using cell proliferation MTS-PMS assay for anti-*T. gondii* activity. To achieve this aim, first the toxicity of the potential compounds towards Vero host cells was tested. Subsequently the non-cytotoxic compounds were evaluated for their efficacy against *T. gondii.* The *T. gondii* infected-Vero cells were treated with various concentrations of the NCI compound. The fluctuations of optical density of each well may be caused by the formation of bubbles resulting from the pipetting action or from the nature of the compounds. To reduce errors caused by the fluctuations, multiple readings of absorbance values were taken every half hour until the end point.

Based on virtual screening, eight NCI compounds i.e., NSC77468, NSC660301, NSC81462, NSC38968, NSC81213, NSC 108972, NSC 153391 and NSC343344 were obtained and further tested for *in-vitro* cytotoxicity and *in-vitro* anti-*T. gondii* activity. 265 All the tested NCI compounds were assumed to target both the enzymes, UPRTase and AK. The activities of the compounds against *T. gondii* were ranked based on the SI values. NSC 77468 gave the highest SI of 25, followed by NSC81462, NSC 660301, NSC81213, NSC38968, NSC108972 with SI values of 12, 6.5. 5.2. 4.5 and 3.9, respectively. No activity was detected with NSC153391 and NSC343344 (Table 3). Both NSC77468 and NSC81462 gave higher SI values compared to clindamycin. Fig 9 shows the high potency of NSC77468 when incubated with *T. gondii*.

**Table 3.** Summary of selectivities of NCI compounds that target UPRTase and AK in the inhibition of the proliferation of Vero cells and *T. gondii*

| Compound | EC<sub>50</sub> against Vero cells (µg/ml) | EC<sub>50</sub> against *T. gondii* | Selectivity Index (SI) |
| --- | --- | --- | --- |
| Clindamycin | 46.63 | 4.270 µg/ml (9.3 nM) | 10.9 |
| 5-Fluorouracil | 9.72 | - | - |
| NSC 77468 | 47.80 | 1.917 µg/ml ( 8.5 nM) | 25 |
| NSC 81462 | 39.01 | 3.240 µg/ml (17.8 nM) | 12.0 |
| NSC 660301 | 22.29 | 3.433 µg/ml (15 nM) | 6.5 |
| NSC 81213 | 24.56 | 4.745 µg/ml (20 nM) | 5.2 |
| NSC 38968 | 21.49 | 4.815 µg/ml (37 nM) | 4.5 |
| NSC 108972 | 28.03 | 7.168 µg/ml (32 nM) | 3.9 |
| NSC 153391 | 24.74 | Inactive | - |
| NSC 343344 | 29.55 | Inactive | - |

**Fig 9.**
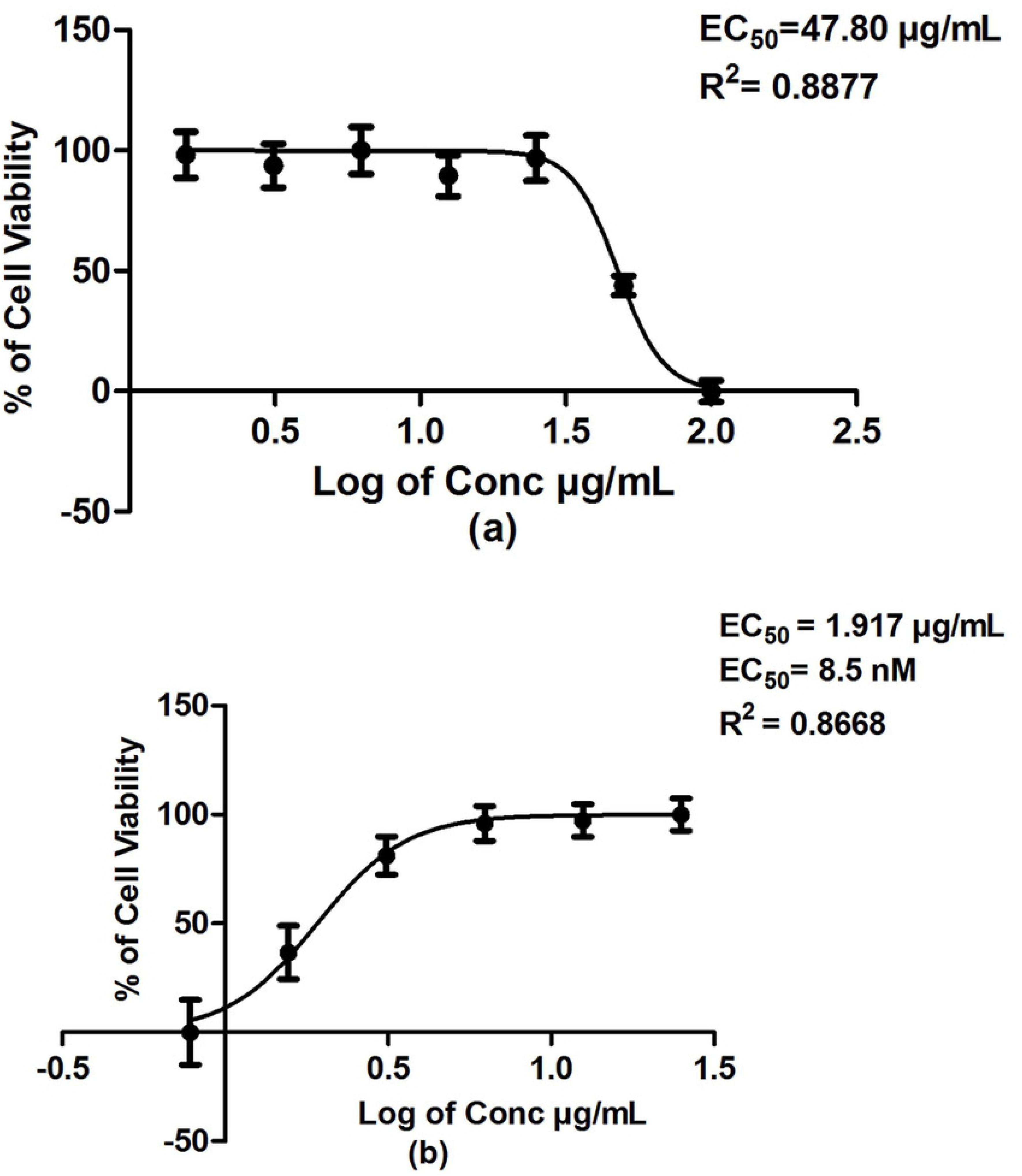
(a) Cytotoxic effect against Vero cells upon increasing NSC77468 concentration and (b) *T. gondii*-infected Vero cells treated with increasing NSC77468 concentration.

Jin et al. (2009) [24] reported novel compounds of (1,5-bis(4-methoxyphenyl)-6,7-dioxabicyclo [3.2.2] nonane) and (1,5-bis(4-fluorophenyl)-6,7-dioxabicyclo[3.2.2] nonane) as having anti-*T. gondii* activities, with SI values of 4.9 and 3.9 respectively. Both compounds were artemisinin derivatives that contain endoperoxide bridge and showed SI values of more than the positive control, spiramycin (SI of 1.0).

NSC77468 showed the highest SI (25) in the present study, demonstrating the effectiveness of the compound against *T. gondii* while being safe to mammalian cells. To the best of our knowledge, this is the first report on the good activity of NSC77468 against *T. gondii.* This compound is relatively rigid, with two rotatable bonds, and has a tendency to be planar, with less chiral centers, and contain pharmacologically desirable features such as no obvious leaving groups, no weakly bonded heteroatoms, no organometallics and no polycyclic aromatic hydrocarbons.

## Conclusions

This study demonstrated that the approach to drug discovery by initially using computer based *in-silico* screening followed by laboratory-based bio-assay can lead to new findings of potential candidates against infectious agents. Virtual screening of dual-target compounds with the combination of potent bioassay activity resulted in identification of compounds that showed potent activities against *T. gondii*. Compared to clindamycin, NSC77468 and NSC81462 gave better EC_50_ and SI values against *T. gondii*. NSC77468 also showed lower toxicity against Vero cells. Thus, NSC77468 could be considered as a potential alternative compound to clindamycin for the treatment of toxoplasmosis.

## Acknowledgments

This study was partially funded by Science Fund from Ministry of Science and Innovation, No. 02-01-05-SF0428 and Universiti Sains Malaysia 1001/PFARMASI/870031. NHS was supported by the Malaysian Institute of Pharmaceuticals and Nutraceuticals through Agilent Bio-analytical Industrial Training Program (BIDP).

## Supporting information

**S1 Fig. The superimposition of docked conformation of 5-fluorouracil (green) and the conformation of crystal structure (blue) in the binding pocket of UPRTase for control docking.**

**S2 Fig. The superimposition of docked conformation of 2,7-iodotubercidin (pink) and the conformation of crystal structure (orange) inthe binding pocket of AK for control docking.**

**S3 Fig. Top eight NCI compounds target to UPRTase and AK.**

**S4 Fig. Hydrogen bond interaction formed between (a) 5-fluorouraciland (b) NSC77468 with ILE229, GLY234 in the active site of UPRTase. Hydrogen bond interaction formed between (c) 2,7-iodotubercidin (blue carbon) and (d) NSC77468(blue carbon) with amino acid residues (white carbon), ACP799 (yellow carbon)in AK active site. Diagram generated from Discovery Studio 4.1.**

**S5 Fig. Interaction formed between 5-Fluorouracil, 2,7-iodotubercidin and NSC77468 with amino acid residues of (a)UPRTase and b) AK. Diagram generated from Discovery Studio 4.1; green residues represent non-polar contact while purple residues represent polar contact.**

**S6 Fig. Aromatic interaction formed between (a) 2,7-iodotubercidin with TYR 169 in the active site of AK while NSC77468 with (b) ARG 136 and (c) TYR 228 in the active site of AK and UPRTase respectively.**

**S7 Fig. The hydrogen bond interaction of NSC77468 (yellow carbon) together with the substrate, ACP (blue carbon) in the binding location of AK.**

**S8 Fig. (a) Cytotoxic effect against Vero cells upon increasing clindamycin con-centration; and (b) *T. gondii*-infected Vero cells treated with increasing clindamycin concentration.**

**S9 Fig. (a) Cytotoxic effect against Vero cells upon increasing NSC77468 concentration and (b) *T. gondii*-infected Vero cells treated with increasing NSC77468 concentration.**

